# Anti- and pro-inflammatory milieu differentially regulate differentiation and immune functions of oligodendrocyte progenitor cells

**DOI:** 10.1101/2023.11.27.568868

**Authors:** Omri Zveik, Ariel Rechtman, Livnat Brill, Adi Vaknin-Dembinsky

**Author notes:** **<u>Corresponding author</u>:** Adi Vaknin-Dembinsky MD, PhD Hadassah Medical Center, Ein–Kerem P.O.B. 12000, Jerusalem 91120, Israel.

## Abstract

Oligodendrocyte progenitor cells (OPC) were regarded for years solely for their regenerative role; however, their immune-modulatory roles have gained much attention recently, particularly in the context of multiple sclerosis (MS). Despite extensive studies on OPCs, there are limited data elucidating the interactions between their intrinsic regenerative and immune functions, as well as their relationship with the inflamed central nervous system (CNS) environment, a key factor in MS pathology. We examined the effects of pro-inflammatory cytokines, represented by interferon (IFN)-γ and tumor necrosis factor (TNF)-α, as well as anti-inflammatory cytokines, represented by interleukin(IL)-4 and IL-10, on OPC differentiation and immune characteristics. Using primary cultures, ELISA, and immunofluorescence stainings, we assessed differentiation capacity, phagocytic activity, major histocompatibility complex (MHC)-II expression, and cytokine secretion. We observed that the anti-inflammatory milieu (IL4 and IL10) reduced both OPC differentiation and immune functions. Conversely, exposure to TNF-α led to intact differentiation, increased phagocytic activity, high levels of MHC-II expression, and cytokines secretion. Those effects were attributed to signaling via TNF-receptor-2 (TNFR2) and counteracted the detrimental effects of IFNγ on OPC differentiation. Our findings suggest that a pro-regenerative, permissive inflammatory environment is needed for OPCs to execute both regenerative and immune-modulatory functions.

## Introduction

Multiple sclerosis (MS) is a chronic autoimmune disorder with a heterogeneous range of clinical manifestations. These manifestations span from a relapsing course (relapsing-MS; rMS) to a progressively deteriorating clinical course (progressive-MS; pMS). The rMS form is characterized by a systemic adaptive autoimmune reaction against the myelin of the central nervous system (CNS), and peripheral immune cell infiltration, resulting in demyelination^1–3^. Conversely, in pMS the inflammation is mainly compartmentalized behind a relatively closed blood-brain barrier^1,2,4^.

Remyelination failure is one of the main obstacles in MS and is considered a major therapeutic challenge^5,6^. Effective remyelination requires oligodendrocyte progenitor cells (OPC) activation, migration into demyelinated lesions, and differentiation into mature myelinating-oligodendrocytes^7^. While mostly successful in rMS, this process is considered a key pathological feature in pMS, primarily due to a differentiation block^6,8–11^. This is evidenced by progenitors in an arrested maturation state in 60-70% of demyelinated pMS lesions, suggesting that OPCs can migrate but fail to differentiate into myelinating oligodendrocytes^12^.

For many years, OPCs were exclusively regarded for their pro-myelinating potential^13,14^. However, the recognition of OPCs’ immune-modulatory properties nowadays is well-established^15–17^. OPC can secrete cytokines and chemokines such as C-C Motif Chemokine Ligand-2 (CCL2) and Tumor necrosis factor(TNF)α in response to injury^11,18^, they can present antigen via major histocompatibility complex (MHC)-I/II^11,17,19^, affect T-cell proliferation^11,17,20^, and phagocytose myelin debris^17^.

While OPCs’ immune functions are now acknowledged, few studies have explored the impact of an inflammatory environment on these functions, with the majority primarily concentrating on their pro-regenerative roles. Evidence indicates OPCs’ pro-regenerative effects in anti-inflammatory settings: interleukin (IL)4 delivery enhanced white-matter integrity post-stroke^21^, and OPCs’ treatment with conditioned media from microglia exposed to IL13 or IL10, but not interferon-γ (IFNγ), significantly enhanced oligodendrocyte differentiation^22^. In contrast, pro-inflammatory conditions, such as IFNγ are known to impede OPC differentiation^23^. TNFα has been linked to OPCs’ differentiation via TNF receptor-2 (TNFR2) signaling^24–26^. Oligodendroglial TNFR2 ablation exacerbated experimental autoimmune encephalomyelitis (EAE), increased axonal and myelin damage, and reduced remyelination^26^. This also influenced OPC immune responses, as seen in gene and cytokine expression changes^25^.

Our previous work indicated that cerebrospinal-fluid (CSF) of pMS patients reduced differentiation and immune functions of OPCs compared to CSF of rMS patients, where these functions remained intact^11^.

Here, we examined how pro- and anti-inflammatory environments affect OPCs’ immune and differentiation capacities. Based on our prior findings, we hypothesized that a pro-inflammatory milieu would enhance differentiation and immune functions compared to an anti-inflammatory one. Characterizing the distinct effects of these environments could deepen our understanding of MS pathology and facilitate the development of novel therapeutic strategies for MS patients.

## Materials and methods

### Approvals

The research reported in this study complied with all relevant ethical regulations for animal testing and research and was approved by the Hebrew University Institutional Animal Care and Use Committee (MD-20-16227-1;MD-22-17079-1).

### Primary OPC cultures

Primary OPC cultures were isolated from naïve P0-1 neonatal C57/BL6 mice cortices as previously described^27^. After shaking, fibroblast growth factor-2 (FGF-2;20ng/mL;R&D systems) and Platelet-derived growth factor (PDGF-AA;20ng/mL;R&D systems) were added to cultures daily unless otherwise specified. Assessments of OPC marker NG2 on day two revealed 91.7±1.6% NG2+ cells.

### Cytokines assays

Mouse primary OPCs were supplemented with PDFG-AA and FGF-2 for the first 48 hours after shaking. Then, media were replaced, and recombinant mouse pro- or anti-inflammatory cytokines were added into the OPC media as specified in each assay. As MS is characterized by simultaneous rise in pro-inflammatory and systemic upregulation of anti-inflammatory cytokines^28^, we selected key cytokines for each. Pro-inflammatory IFNγ(Cat#315-05,PeproTech, 10ng/mL) and TNFα (Cat#315-01A,PeproTech,200ng/mL), were chosen for their pivotal roles in MS relapses and progression, as well as due to their prevalence in MS patients’ CSF^3,29–31^. IL4(Cat#214-14,PeproTech,10ng/mL) and IL10(Cat#210-10,PeproTech,100ng/mL) were chosen to represent anti-inflammatory environment in MS^28,32^.

### Differentiation assays

48 hours after shaking, media were replaced, and combinations of Triiodothyronine(T3;60ng/mL;Sigma-Aldrich) and pro- or anti-inflammatory cytokines were added into the OPC media for seven days.

### MHC assays

Myelin oligodendrocyte glycoprotein(MOG)-35-55 peptide(Sigma-Aldrich,100µg/ml), IFNγ(10ng/ml;Cat#315-05,PeproTech) and pro- or anti-inflammatory cytokines were supplemented into mouse primary OPC media for 72 hours to allow transcription and translation of MHC-II pathway mediators and permit for expression time, as described previously^11^.

### Latex beads phagocytosis

OPCs were stimulated using IFNγ(10ng/ml) in a combination of pro- and anti-inflammatory cytokines for 24 hours. Then, latex beads were added for two hours(Cat#L3030-1ML,3µl/1mL,Sigma-Aldrich). Following two hours, cells were immune stained for NG2 marker. Uptake analysis was quantified manually in a blinded manner.

### TNFR2 neutralization assays

Mouse primary OPCs were treated with neutralizing antibody against TNFR2(anti-TNFR2; Cat#50128-RN204,RRID:AB_2860178,Sino Biological,6ug/mL) for 1 hour prior to exposure to pro-inflammatory cytokines, as previously described^33^. PBS served as control.

### Immunostaining

For cell surface markers, staining was performed on living cells followed by fixation^34^. Anti-NG2(Cat#AB5320, RRID:AB_91789, Millipore, 1:100) was used to identify OPCs, anti-O1(Cat#MAB1327, RRID:AB_357618, R&D System, 1:100) for mature oligodendrocytes, and anti-IBL-5/22(Cat#sc-59,322, RRID:AB_831551, Santa Cruz Biotechnology, 1:100) for evaluation of MHC-II expression. Goat anti-rabbit Alexa Fluor-488(Cat#A-11034, RRID:AB_2576217, Invitrogen, 1:200), goat anti-mouse Alexa Fluor-488(Cat#A-11001, RRID:AB_2534069, Invitrogen, 1:200), and goat anti-rat Alexa Fluor-555(Cat#A-21434, RRID:AB_141733, Invitrogen, 1:200) were used as secondary antibodies appropriately. Nuclei were counterstained with 4′,6-diamidino-2-phenylindole(DAPI; Cat#H-1200, RRID:AB_2336790, Vector Laboratories). Quantification was performed using ImageJ software(NIH,https://imagej.nih.gov/ij,RRID:SCR_003070) by measuring positively stained cells relative to total DAPI. Quantifications are represented as mean percentages from total DAPI+ cells ±SD and are from at least 15 random fields captured in 3 or more independent experiments.

### Assessment of oligodendrocyte morphological complexity

Fractal dimension(FD) analysis was performed to evaluate morphological complexity of oligodendrocytes following differentiation essays, as described previously^11^. A numerical value close to 1 represents cells with low morphological complexity. This procedure was performed for >25 cells per condition for each independent experiment.

### Protein concentration measurements

Enzyme-linked immunosorbent-assay (ELISA) was used to measure proteins secretion in culture superannuants following the manufacturer’s protocol. The following proteins were measured: CCL2(Cat#DY479-05, R&D System,), IL1b(Cat#DY401-05,R&D System), IL6(Cat#431304, BioLegend; a generous gift of Prof. Talma Brenner), and IL10(Cat#431414, BioLegend; a generous gift of Prof. Talma Brenner). IL1b levels were undetected among all groups.

### RNA isolation and reverse transcription

OPCs were incubated with T3 and pro-inflammatory cytokines for 72h. RNA was extracted using Tri-reagent and cDNA was synthesized^34^. Quantitative polymerase chain reaction(PCR) was conducted with PerfeCTa SYBR-Green FastMix-Rox(Quanta-Biosciences) and the StepOnePlus rt-PCR system(Applied Biosystems). We used the 2-ΔCT method for statistical analysis, normalizing all target mRNAs to HPRT reference gene.

#### Primers(Agentek)

HPRT:F:5‘CATGGACTGATTATGGACGGAC R:5‘ACAGAGGGCCACAATGTGATG

CCL2:F:5‘GGCTCAGCCAGATGCAGT R:5‘GCTCTCCAGCCTACTCATTGG

IL1β:F:5‘CCAAGCITCCTTGTGCAAGIGT R:5‘GGTGGCATTTCACAGTIGAGTTC

IL10:F:5‘CAGCCAGGTGAAGACITICITIC R:5‘CTGCATTAAGGAGTCGTTAGCA

### Statistical analyses

We assessed distribution using the Shapiro–Wilk test. Given the normal distribution, we applied one-way ANOVA with Šídák’s post hoc test and two-way ANOVA with Tukey’s post hoc test for multiple comparisons. Each figure legend specifies the tests used and their significance levels. Data are presented as mean±SD, or as detailed in each figure. Error bars denote standard deviation. Each assay was repeated independently at least three times.

## Results

### 1. OPC differentiation is impeded by anti-inflammatory cytokines and IFNγ, while preserved by TNFα

Initially, we studied the effect of pro- and anti-inflammatory mediators on OPCs’ ability to differentiate. OPCs were cultured with T3(60ng/mL)^35^ and pro- or anti-inflammatory cytokines for seven days, followed by immunofluorescence staining^36^ (Figure 1a-b). IL4 and IL10, representing anti-inflammatory mediators, significantly decreased OPC differentiation into mature oligodendrocytes (26.39±3.32% and 24.15±3.48%, respectively, Figure 1b-c), compared to T3 alone (63.1±4.64%,p<0.0001 and p<0.0001,respectively,Figure 1b-c). A similar inhibitory effect was observed upon exposure to IFNγ, a pro-inflammatory mediator (25.74±3.37%, Figure 1b-c). The exposure to TNFα led to intact differentiation levels (60.9±4.63%) compared to T3 alone (p=0.0634,Figure 1b-c). Moreover, exposure of OPC to both TNFα and IFNγ together led to higher levels of differentiation (55.32±4.19%) compared to exposure to IFNγ alone (p<0.0001).

**Figure 1.**
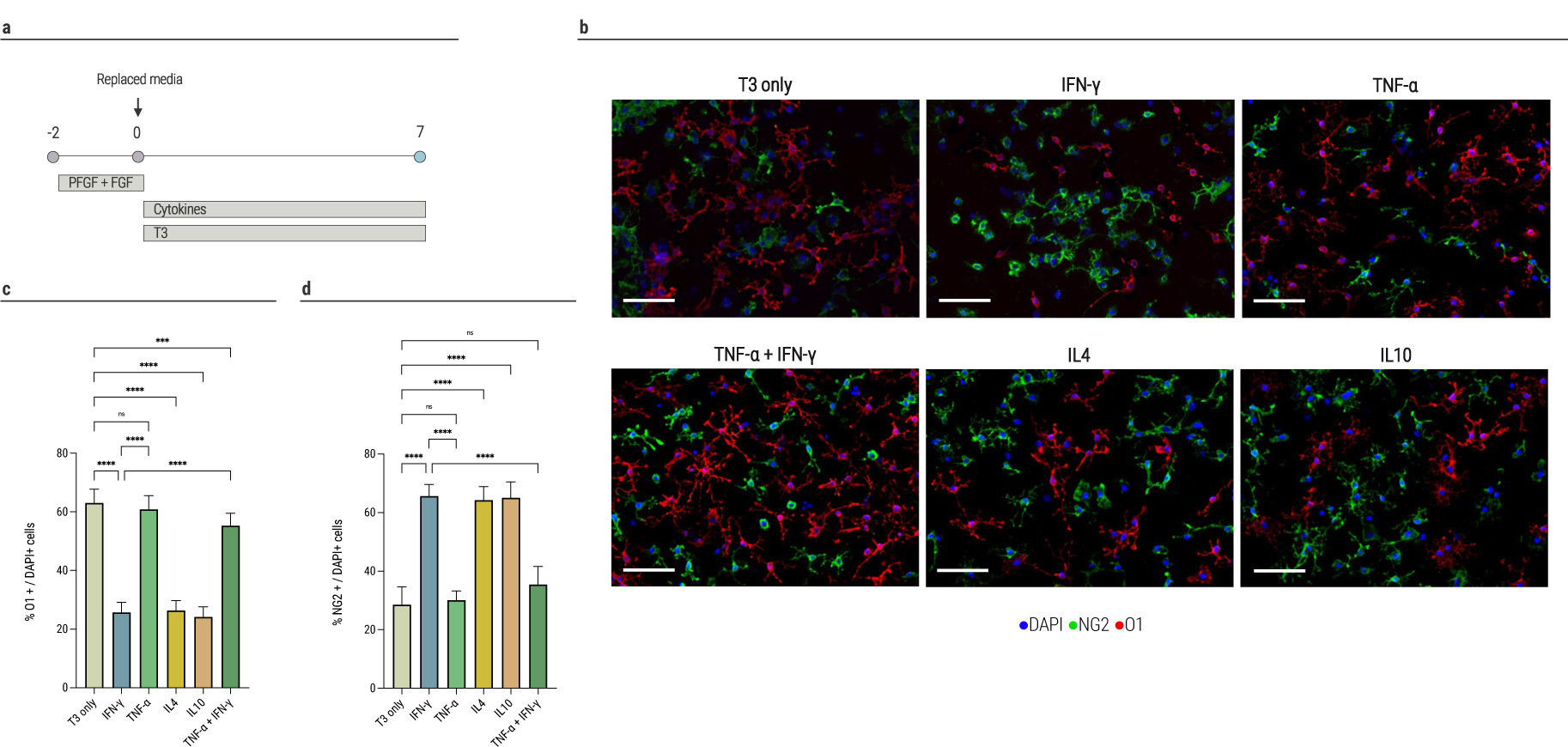
OPC differentiation is impeded by anti-inflammatory cytokines and IFNγ, while preserved by TNFα. **(a)** Time-course of differentiation experiments. OPCs were supplemented with PDGF and FGF2 for 48h. Then, each culture was exposed to simultaneously to thyroid hormone(T3) and either anti- or pro-inflammatory cytokines for 7 days and evaluated using OPC marker NG2 and mature oligodendrocyte marker O1,(**b)** Representative immunofluorescence analysis of mature oligodendrocyte marker O1 and progenitor marker NG2 on day 7(scale bar = 50 μm),(**c)** Exposure to IFNγ(25.74±3.37%) or anti-inflammatory cytokines(IL4: 26.39±3.32%; IL10: 24.15±3.48%) significantly decreased OPC differentiation into mature oligodendrocytes as compared to T3 alone(63.1±4.64%), TNFα(60.9±4.63%), or TNFα and IFNγ together(55.32±4.19%); n=9 for each group,(**d)** OPCs exposed to IFNγ(65.63±3.96%) or anti-inflammatory cytokines(IL4: 64.24±4.63%; IL10: 65.07±5.38%) expressed higher levels of progenitor marker NG2 after 7 days in culture in comparison to T3 alone(28.55±6.09%), TNFα(30.13±3.1%), or TNFα and IFNγ together(35.43±6.18%); n=5 for each group. Significance was determined by one-way ANOVA analysis followed by Šídák’s multiple comparison analysis (p * ≤ 0.05, ** ≤ 0.01, *** ≤ 0.001, **** ≤ 0.0001, ns = not significant). Data are presented as means±SD. Each assay was repeated independently at least three times.

Analysis for NG2 expression, an OPC marker, revealed significantly higher expression upon exposure to IFNγ (65.63±3.96%), IL4 (64.24±4.63%), or IL10 (65.07±5.38%), compared with OPCs exposed to T3 alone, TNFα, or TNFα and IFNγ together (28.55±6.09%, 30.13±3.1%, and 35.43±6.18, respectively, p<0.0001, Figure 1d).

Our findings imply that an anti-inflammatory milieu might hinder OPC differentiation capabilities and impair the remyelination process in the inflamed-CNS. Additionally, TNFα seems to preserve OPC differentiation and may play an essential role in the delicate interplay between immunological function and regenerative abilities of OPCs in MS.

#### 1.1. Lower morphological complexity of O1+ oligodendrocytes upon culture with pro-inflammatory cytokines

It has been suggested that mature oligodendrocytes have more complex branches^36^. Alongside, our prior research indicates a negative correlation between oligodendrocytes’ immune activation levels and the complexity of their branches^11,27^, as supported by earlier observations^37,38^.

Using fractal dimension and skeleton analyses, we performed combined quantification of the morphological pattern of O1+ oligodendrocytes upon culture with pro- or anti-inflammatory mediators for seven days. Oligodendrocytes cultured with pro-inflammatory mediators (TNFα, IFNγ) had lower morphological complexity, represented by a significantly lower FD index, than OPCs exposed to anti-inflammatory mediators (IL4,IL10,Figure 2a-g). Oligodendrocytes exposed to pro-inflammatory cytokines also had lower end points/cell index and branch length/cell compared to those exposed to anti-inflammatory cytokines (Figure 2h-i). Remarkably, simultaneous TNFα and IFNγ exposure resulted in higher levels of end points/cell index and branch length/cell compared to their separate applications (Figure 2b,c,f,h,i).

**Figure 2.**
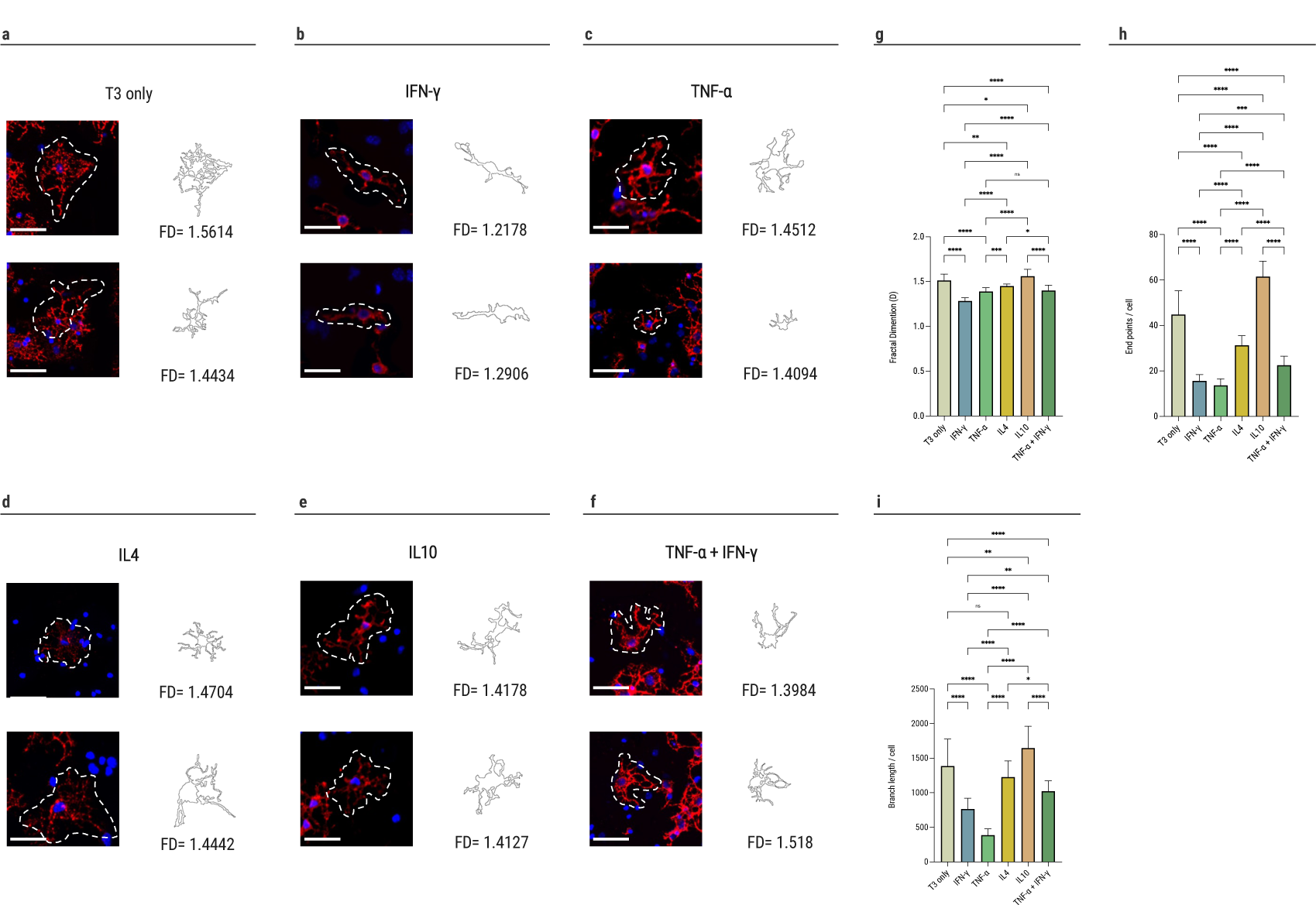
Lower morphological complexity of O1+ oligodendrocytes upon culture with pro-inflammatory cytokines. (**a-f)** Representative images of O1+ oligodendrocytes and cell outline with the appropriate fractal dimension(FD) value(scale bar = 30 μm),**(g)** Higher levels of morphological complexity among cells exposed to anti-inflammatory cytokines(IL4: 1.45±0.02; IL10: 1.56±0.08) or T3(1.51±0.07) compared with IFNγ(1.29±0.04), TNFα(1.39±0.04), or TNFα and IFNγ together(1.4±0.06),(**h)** Skeleton analysis resulted in higher endpoints/cell among cells exposed to anti-inflammatory cytokines(IL4: 31.32±4.21; IL10: 61.52±6.74) or T3(44.8±10.41) compared with IFNγ(15.6±2.8), TNFα(13.689±2.81), or TNFα and IFNγ together(22.5±3.88),(**i)** Skeleton analysis resulted in higher branches length/cell among cells exposed to anti-inflammatory cytokines(IL4: 1226.0±233.7; IL10: 1646.0±315.4) or T3(1386.0±389.4) compared with IFNγ(766.4±153.8), TNFα(385.6±93.03), or TNFα and IFNγ together(1023.0±149.5). Significance was determined by one-way ANOVA analysis followed by Šídák’s multiple comparison analysis (p * ≤ 0.05, ** ≤ 0.01, *** ≤ 0.001, **** ≤ 0.0001, ns = not significant). Data are presented as means±SD. Each assay was repeated independently at least three times.

### 2. Anti-inflammatory cytokines impede OPC phagocytic capability, MHC-II expression, and cytokines secretion

#### 2.1. Anti-inflammatory cytokines impede OPC phagocytic capability

The CNS environment in MS contains inflammatory mediators and myelin degradation products, which are known to influence MS inflammatory cortical activity, activation of CNS innate-immune cells, and OPC remyelination ability. Immune cells, primarily microglia, are responsible for clearing myelin debris, which contains proteins that inhibit OPC differentiation. Our prior study demonstrated that OPC exposed to CSF from rMS or pMS patients differ mainly in pathways involved in immune response, specifically phagocytosis^11^. This is further substantiated by a study demonstrating the phagocytic abilities of OPCs^17^. In this regard, we decided to investigate whether and which anti- or pro-inflammatory cytokines could affect this ability.

OPCs were initially stimulated with IFNγ (10ng/mL), then pro-inflammatory(TNFα) or anti-inflammatory cytokines (IL4,IL10) were added for 24h. Afterward, latex beads were added for 2h, followed by immunofluorescence staining (Figure 3a). Simultaneous exposure of OPCs to TNFα and IFNγ significantly enhanced their phagocytic ability (36.29±2.81%) compared to IFNγ alone (28.89±2.86%), IL4 and IFNγ(23.62±2.51%), IL10 and IFNγ (17.37±2.14%), or both IFNγ, IL4, and IL10 (24.5±2.83%, p<0.0001, Figure 3b-c).

**Figure 3.**
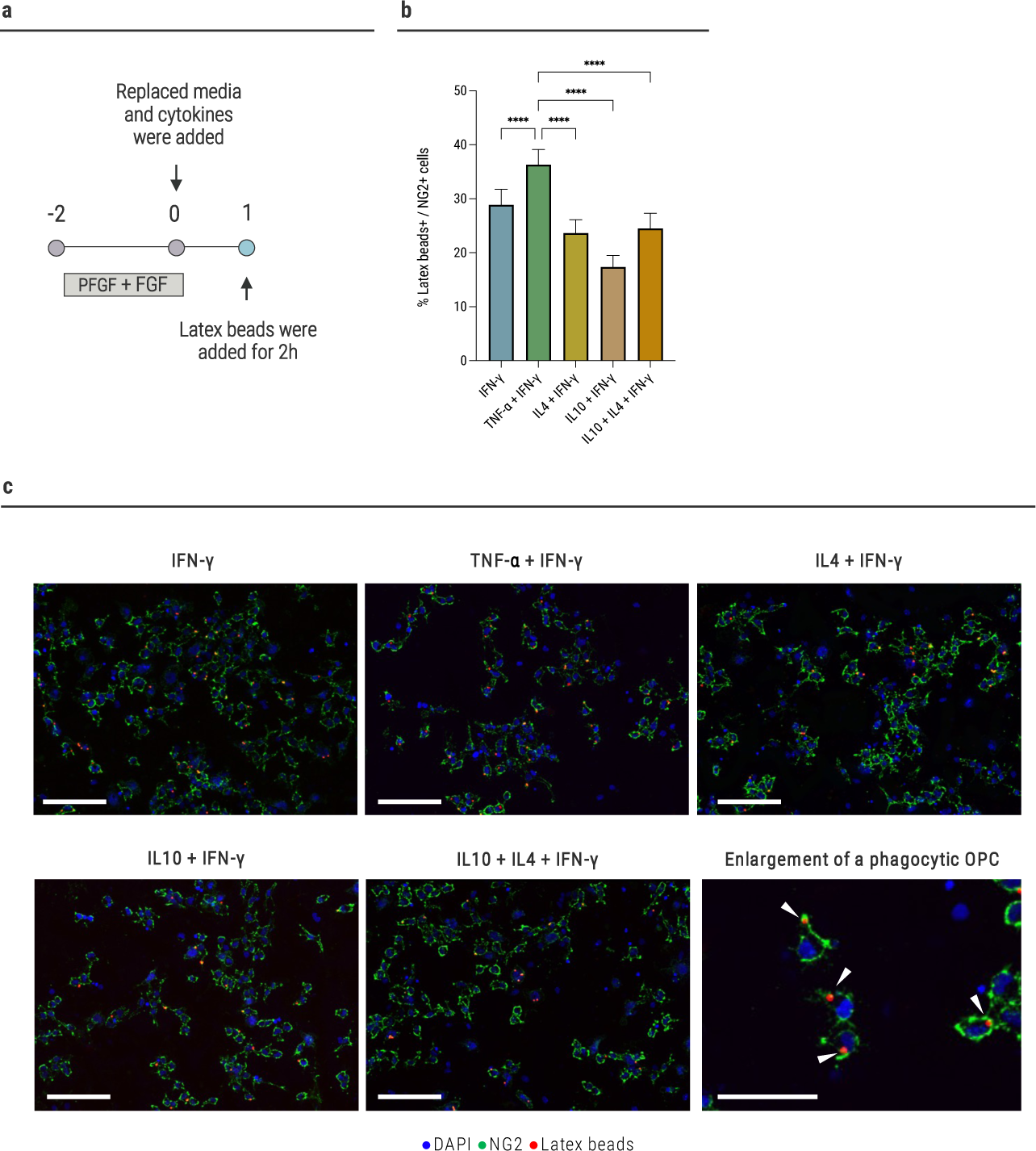
Anti-inflammatory cytokines impede OPC phagocytic capability. **(a)** Time-course experiments of assessment of phagocytic capability. Primary OPCs were cultured under PDGF and FGF conditions(daily dose of 20 ng/mL) for 48h. After 48h, OPCs were stimulated with IFNγ(10ng/mL), then pro-inflammatory(TNFα) or anti-inflammatory cytokines(IL4 and IL10) were added for 24h. Afterward, latex beads were added to the cultures for 2h, followed by evaluation of phagocytosed latex beads out of NG2+ cells using immunofluorescence staining., **(b)** Simultaneous exposure of OPCs to TNFα and IFNγ significantly enhanced their phagocytic ability(36.29±2.81%) compared to IFNγ alone(28.89±2.86%), IL4(23.62±2.51%), IL10(17.37±2.14%), or both IL4 and IL10(24.5±2.83%); n=7 for each group,(**c)** Representative immunofluorescence analysis of NG2+ OPCs and phagocytosed latex beads upon culture with either anti- or pro-inflammatory cytokines for 24h(scale bar = 40 μm). Significance was determined by one-way ANOVA analysis followed by Šídák’s multiple comparison analysis (p * ≤ 0.05, ** ≤ 0.01, *** ≤ 0.001, **** ≤ 0.0001, ns = not significant). Error bars in all graphs represent SD. Each assay was repeated independently at least three times.

These results indicate that TNFα amplifies OPC-mediated clearance of myelin debris more than anti-inflammatory settings.

#### 2.2. Reduced MHC-II expression in OPC exposed to anti-inflammatory cytokines

Antigen presentation is crucial for T-cell response. As we previously demonstrated that MHC-II expression in OPC is reduced upon exposure to CSF from pMS compared to rMS patients^11^, we aimed to examine the influence of pro- and anti-inflammatory cytokines on OPCs’ ability to express MHC-II.

OPCs were cultured with MOG peptide (100µg/mL) and stimulated with IFNγ (10ng/mL). Additionally, cells were simultaneously exposed to pro-inflammatory (TNFα) or anti-inflammatory (IL4,IL10) cytokines for 72h. We then assessed MHC-II expression using immunofluorescence staining (Figure 4a).

**Figure 4.**
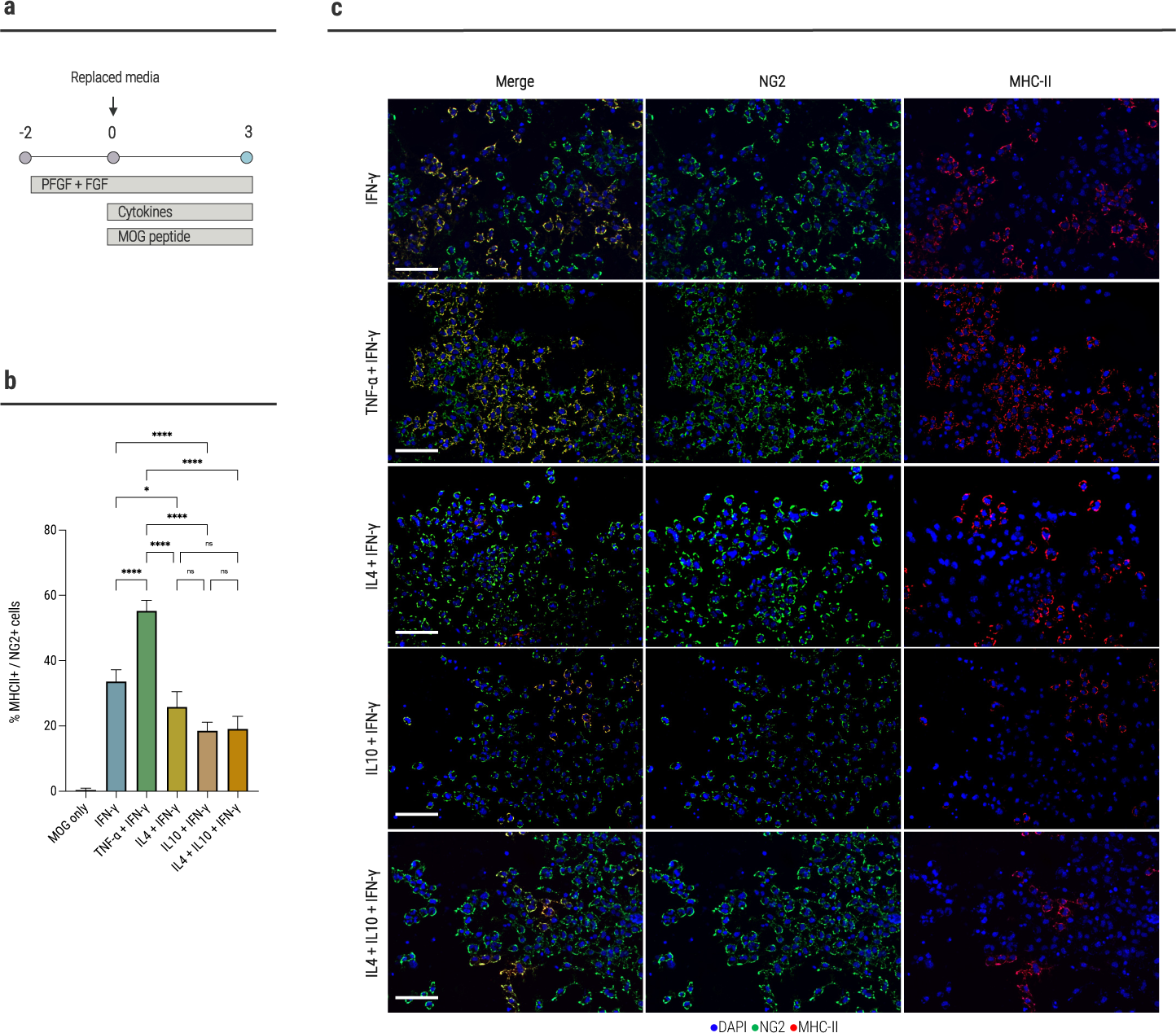
Reduced MHC-II expression in OPC exposed to anti-inflammatory cytokines. **(a)** Time-course experiments of MHC-II expression. Primary OPCs were cultured under PDGF and FGF conditions(daily dose of 20 ng/mL) for 48h. After 48h, OPCs were stimulated with IFNγ(10ng/ml), then pro-inflammatory(TNFα) or anti-inflammatory cytokines(IL4 and IL10) were added for 72h, followed by evaluation of MHC-II expression out of NG2+ cells using immunofluorescence staining,(**b)** Exposure of OPCs to anti-inflammatory cytokines significantly reduced MHC-II expression(IL4: 25.8±4.66%; IL10: 18.47±2.65%; IL4 and IL10 together: 19.12±3.78%) compared to the exposure to IFNγ alone(33.63±3.6%) or TNFα and IFNγ together(55.23±3.22%); n=5 for each group,(**c)** Representative immunofluorescence analysis of MHC-II expression among OPCs upon culture with either anti- or pro-inflammatory cytokines for 72h(scale bar = 40 μm). Significance was determined by one-way ANOVA analysis followed by Šídák’s multiple comparison analysis (p * ≤ 0.05, ** ≤ 0.01, *** ≤ 0.001, **** ≤ 0.0001, ns = not significant). Error bars in all graphs represent SD. Each assay was repeated independently at least three times.

Exposure of OPCs to anti-inflammatory cytokines (IL4 and IL10) significantly reduced MHC-II expression (25.8±4.66% and 18.47±2.65%, respectively) compared to the exposure to IFNγ alone(33.63±3.6%, p=0.0395 and p<0.0001, respectively, Figure 4b-c). Simultaneous IL4 and IL10 exposure showed no synergistic effect (Figure 4b). Notably, the simultaneous exposure of OPCs to two major pro-inflammatory mediators, TNFα and IFNγ, significantly enhanced MHC-II expression (55.23±3.22%) compared to IFNγ alone, IL4, or IL10 (p<0.0001, Figure 4b-c).

#### 2.3. Increased immune gene expression upon exposure of OPCs to pro-inflammatory cytokines

To investigate how a pro-inflammatory environment affects immune gene expression in OPCs, we explored the expression of CCL2, IL1β, and IL10 in OPC differentiation cultures. OPCs were cultured with T3 (60ng/ml) and pro-inflammatory cytokines for 72h, followed by RNA extraction and rt-PCR analysis.

Post-injury CCL2 secretion by OPCs promotes mobilization, inflammation, and regeneration^18^. OPCs exposed to TNFα showed a significant increase in CCL2 gene expression (13.04±3.89) compared to T3 alone (1.16±0.39, p=0.0129). TNFα and IFNγ exposure further elevated CCL2 expression (36.59±7.92) compared to TNFα alone (p<0.0001, Figure 5a).

**Figure 5.**
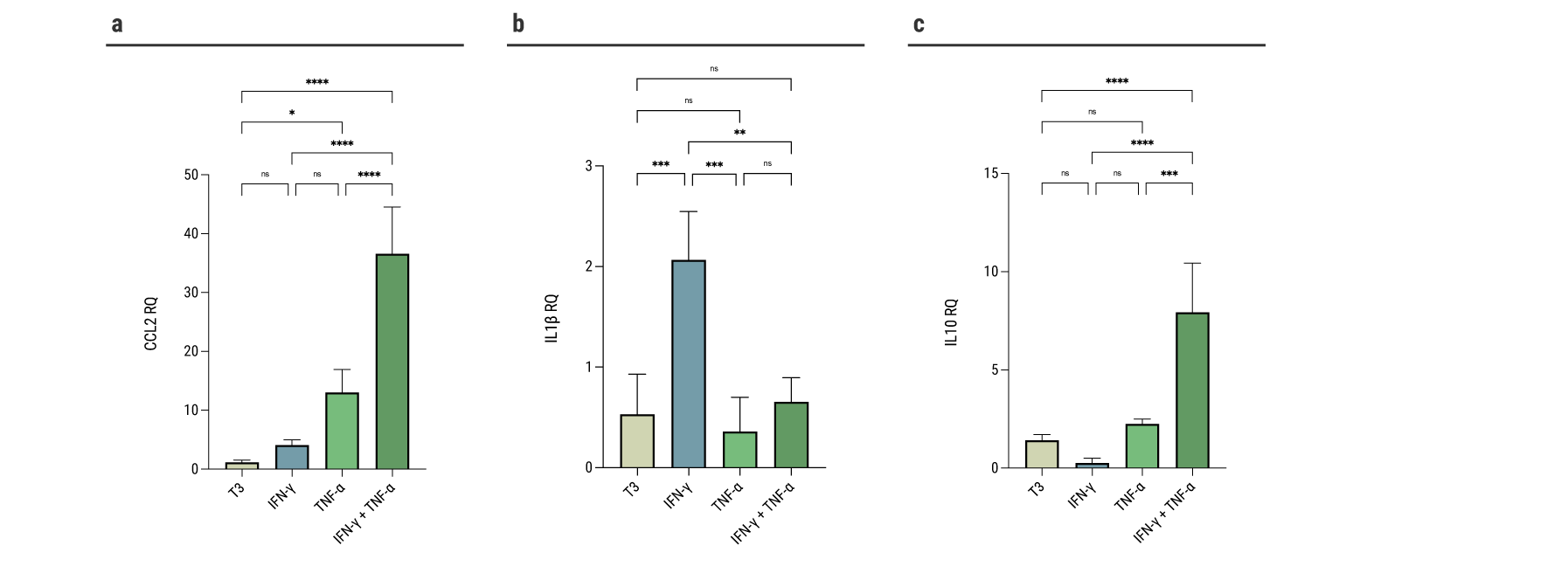
Increased immune gene expression upon exposure of OPCs to pro-inflammatory cytokines. Using rt-PCR analyses, we studied the immune gene expression (CCL2, IL1β, and IL10) of OPCs following culture with T3 and pro-inflammatory cytokines for 72h. (**a)** CCL2: T3 alone: 1.16±0.39; IFNγ: 4.11±0.85; TNFα: 13.04±3.89; IFNγ and TNFα: 36.59±7.92; n=4 for each group,(**b)** IL1β: CCL2: T3 alone: 0.53±0.39; IFNγ: 2.06±0.48; TNFα: 0.35±0.34; IFNγ and TNFα: 0.65±0.23; n=4 for each group,(**c)** IL10: CCL2: T3 alone: 1.41±0.28; IFNγ: 0.25±0.23; TNFα: 2.25±0.23; IFNγ and TNFα: 7.93±2.5; n=4 for each group. Significance was determined by one-way ANOVA analysis followed by Šídák’s multiple comparison analysis (p * ≤ 0.05, ** ≤ 0.01, *** ≤ 0.001, **** ≤ 0.0001, ns = not significant). Error bars in all graphs represent SD. Each assay was repeated independently at least three times.

IL1β, crucial in CNS inflammation and regeneration^18,39^, was also upregulated in OPCs cultured with pro-inflammatory cytokines. IFNγ exposure significantly increased IL1β expression levels (2.06±0.48), surpassing those seen with T3 alone(0.53±0.39,p=0.0004), TNFα (0.35±0.34, p=0.0002), or TNFα and IFNγ (0.65±0.23, p=0.0009, Figure 5b).

Finally, IL10 gene expression was highest with combined TNFα and IFNγ exposure (7.93±2.5), significantly exceeding levels with IFNγ (0.25±0.23, p<0.0001), TNFα (2.25±0.23, p=0.0002), or T3 alone (1.41±0.28, p<0.0001, Figure 5c).

#### 2.4. Increased secretion of CCL2, IL6, and IL10 upon exposure of OPCs to TNFα

To better understand the effect of the environment on OPCs’ immune phenotype, we analyzed CCL2, IL6, and IL10 secretion in OPCs’ differentiation cultures exposed to pro- or anti-inflammatory cytokines for seven days (Figure 1a).

Exposure of OPCs to TNFα led to higher levels of CCL2(953.8±87.47pg/mL) compared to T3 alone (302.8±86.33pg/mL, p<0.0001), IFNγ (799.2±47.00pg/mL, p=0.0281), IL10 (606.7±154.0pg/mL, p<0.0001), or IL4 (477.9±115.1pg/mL, p<0.0001, Figure 6a).

**Figure 6.**
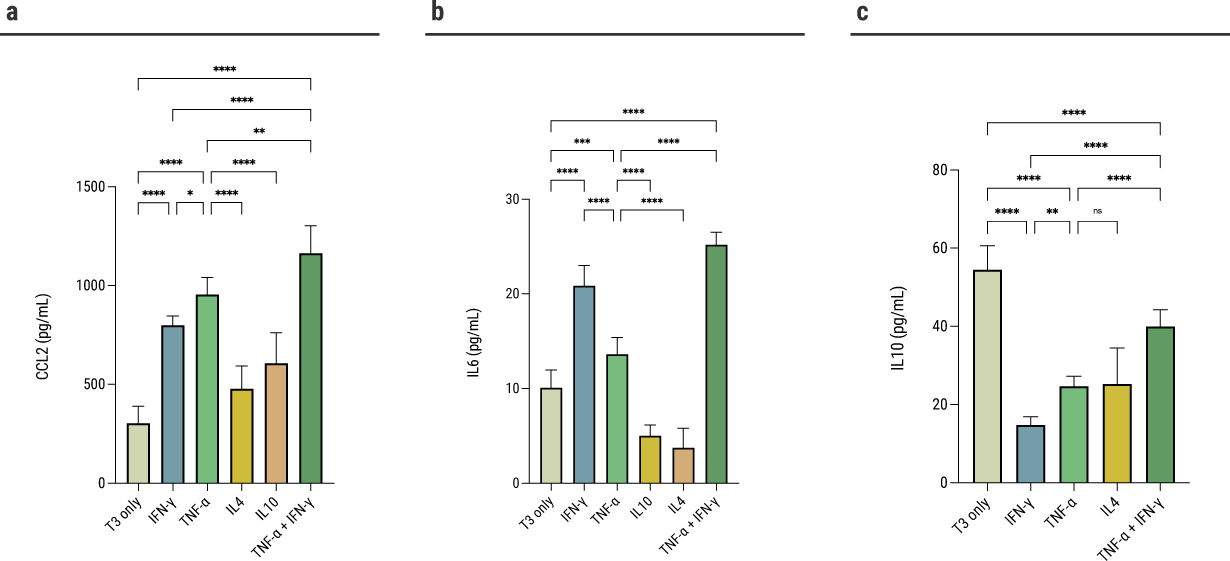
Increased secretion of CCL2, IL6, and IL10 upon exposure of OPCs to TNFα. Using ELIAS analyses, we studied the secretion of CCL2, IL6, and IL10 in the supernatant of OPCs following culture with T3 and either anti- or pro-inflammatory cytokines for 7 days.(**a)** CCL2: T3 alone: 302.8±86.33pg/mL; IFNγ: 799.2±47.0pg/mL; TNFα: 953.8±87.47pg/mL; IL4: 477.9±115.1pg/mL; IL10: 606.7±154.0pg/mL; IFNγ and TNFα: 1163.0±139.5pg/mL; n=9 for each group,(**b)** IL6: T3 alone: 10.09±1.88pg/mL; IFNγ: 20.87±2.13pg/mL; TNFα: 13.63±1.76pg/mL; IL4: 3.75±2.07pg/mL; IL10: 5.02±1.13pg/mL; IFNγ and TNFα: 25.19±1.33pg/mL; n=9 for each group,(**c)** IL10: T3 alone: 54.5±6.14pg/mL; IFNγ: 14.81±2.07pg/mL; TNFα: 24.69±2.55pg/mL; IL4: 25.27±9.2pg/mL; IFNγ and TNFα: 39.97±4.3pg/mL; n=9 for each group. Significance was determined by one-way ANOVA analysis followed by Šídák’s multiple comparison analysis (p * ≤ 0.05, ** ≤ 0.01, *** ≤ 0.001, **** ≤ 0.0001, ns = not significant). Error bars in all graphs represent SD. Each assay was repeated independently at least three times.

Simultaneous TNFα and IFNγ exposure led to higher CCL2 secretion levels (1163.0±139.5pg/mL) compared to TNFα or IFNγ alone (p=0.0012 and p<0.0001, respectively, Figure 6a).

The pro-inflammatory cytokine IL6 is known for its harmful effect in the inflamed-CNS. The highest IL6 levels were measured in cultures exposed simultaneously to TNFα and IFNγ (25.19±1.33pg/mL, Figure 6b). The exposure to TNFα alone led to lower levels of IL6 secretion(13.63±1.76pg/mL) compared to IFNγ (20.87±2.13pg/mL, p<0.0001) but higher compared to the exposure to IL4, IL10, or T3 alone (3.75±2.07pg/mL, 5.02±1.13pg/mL, and 10.09±1.88pg/mL, respectively, p<0.0001, p<0.0001, and p<0.001, respectively, Figure 6b).

Last, the highest IL10 levels were measured in supernatants of T3 alone cultures (54.5±6.14pg/mL, Figure 6c). The exposure to TNFα led to higher IL10 secretion levels (24.69±2.55pg/mL) compared to IFNγ (14.81±2.07pg/mL, p=0.0015). Simultaneous exposure to TNFα and IFNγ led to higher IL10 secretion levels (39.97±4.23pg/mL) compared to the exposure to TNFα or IFNγ alone (p<0.0001, Figure 6c).

These results suggest that TNFα potentially activates OPCs, promoting neuroprotection by enhancing the regenerative capabilities of OPCs and attracting immune cells to the lesion site.

### 3. TNFα-mediated differentiation, but not MHC-II expression, is dependent on TNFR2

TNFα, a multifaceted cytokine, is abundantly present in the serum, CSF, and active lesions of MS patients^29,30^. Signaling can occur through two receptors: TNFR1, which primarily promotes neurotoxicity, and TNFR2, which promotes neuroprotection and reparative effects^40^. TNFR2 plays a role in OPC differentiation and has been the focus of research in recent years^24–26^. To study the effects of TNFR2 on OPCs’ immune and differentiation capabilities *in-vitro*, we used a neutralizing antibody against TNFR2.

#### 3.1. TNFR2 neutralization impairs TNFα-mediated OPC differentiation

We aimed to examine whether TNFR2 plays a role in the differentiation observed following TNFα exposure. OPCs were pre-treated with a neutralizing antibody directed to TNFR2 or PBS for 1h. Then, cultures were treated with T3^35^ and pro-inflammatory cytokines for seven days.

In the anti-TNFR2 group, TNFα alone and simultaneous TNFα and IFNγ exposure resulted in significantly lower differentiation levels compared to the control(TNFα alone: 63.02±1.92% vs. 31.39±0.37%, p<0.0001; TNFα and IFNγ: 57.16±5.64% vs. 29.79±2.64%, p<0.0001, Figure 7a,b). No significant difference in O1+ cell percentages was observed in cultures exposed to IFNγ or T3 alone between the anti-TNFR2 and control groups (Figure 7a,b), indicating that T3-induced differentiation is TNFR2-independent. The anti-TNFR2 group showed markedly lower differentiation levels when exposed to TNFα alone or TNFα with IFNγ compared to T3 alone, unlike the similar levels observed without TNFR2 neutralization (p<0.0001, Figure 1c, 6b). Analysis for NG2 expression revealed significantly higher NG2 levels among OPCs cultured with TNFα alone or TNFα together with IFNγ, which were pre-treated with anti-TNFR2 (Figure 7c).

**Figure 7.**
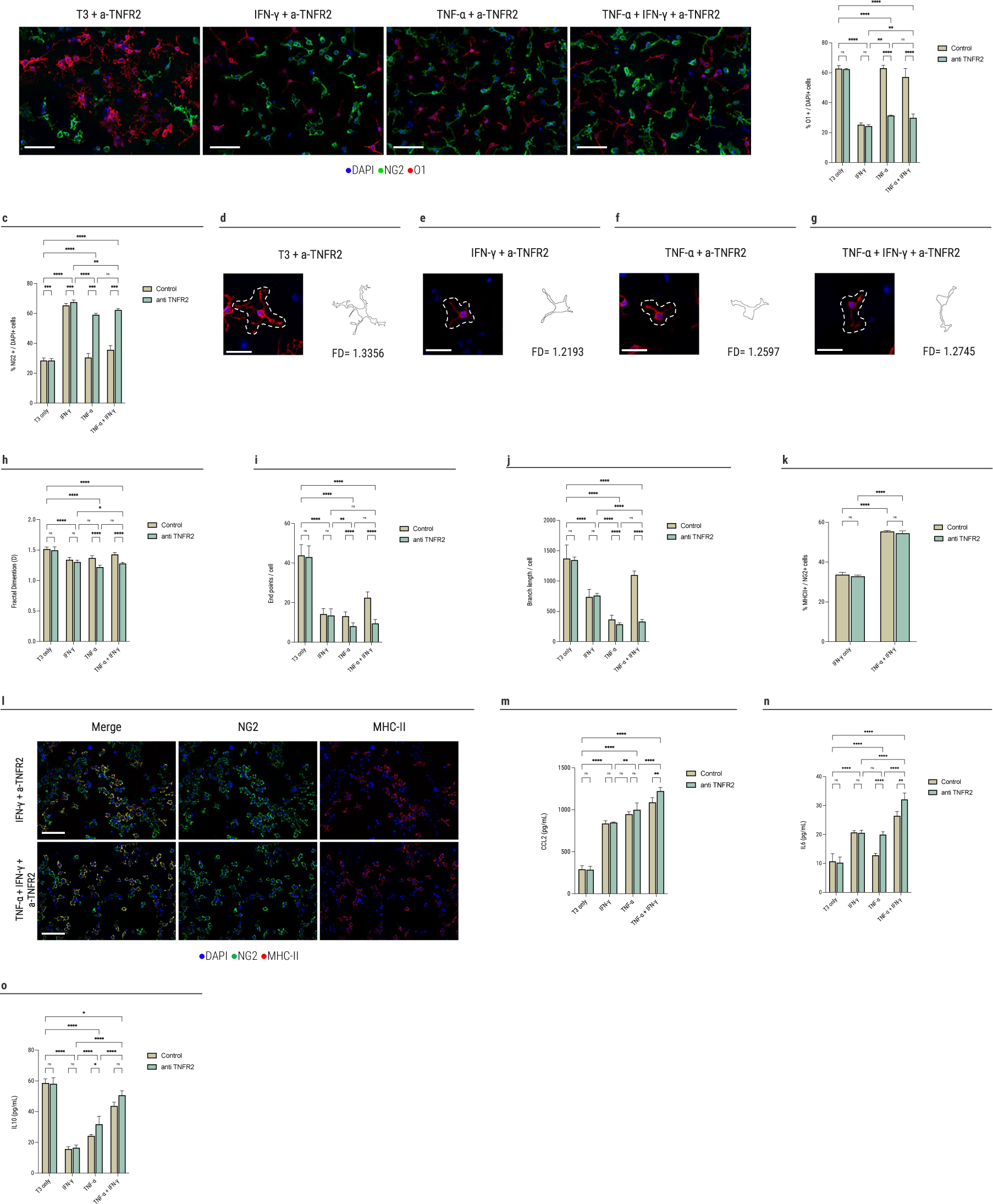
TNFα-mediated differentiation, but not MHC-II expression, is dependent on TNFR2. **(a)** Representative immunofluorescence analysis of mature oligodendrocyte marker O1 and progenitor marker NG2 on day 7 for TNFR2 neutralizations differentiation experiments(scale bar = 50 μm),(**b)** Percentage of O1+ cells out of DAPI on day 7: T3 only: control 62.8±2.0%, a-TNFR2 62.4±0.5%; IFNγ: control 25.25±1.22%, a-TNFR2 24.31±0.9%; TNFα: control 63.02±1.92%, a-TNFR2 31.39±0.37%; IFNγ and TNFα: control 57.16±5.64%, a-TNFR2 29.79±2.64%; n=4 for each group,(**c)** Percentage of NG2+ cells out of DAPI on day 7: T3 only: control 28.55±1.67%, a-TNFR2 28.5±1.34%; IFNγ: control 65.36±1.49%, a-TNFR2 67.54±1.5%; TNFα: control 30.58±2.7%, a-TNFR2 59.1±1.0%; IFNγ and TNFα: control 35.7±2.76%, a-TNFR2 62.39±0.95%; n=4 for each group,(**d-g)** Representative images of O1+ oligodendrocytes and cell outline with the appropriate fractal dimension(FD) value for TNFR2 neutralizations differentiation experiments(scale bar = 30 μm),(**h)** FD index for TNFR2 neutralizations differentiation experiments: T3 only: control 1.52±0.03, a-TNFR2 1.5±0.06; IFNγ: control 1.34±0.04, a-TNFR2 1.31±0.03; TNFα: control 1.37±0.04, a-TNFR2 1.22±0.03; IFNγ and TNFα: control 1.43±0.04, a-TNFR2 1.28±0.02,(**i)** End points/cell index for TNFR2 neutralizations differentiation experiments: T3 only: control 43.8±5.5, a-TNFR2 42.96±5.72; IFNγ: control 14.2±2.84, a-TNFR2 13.46±3.45; TNFα: control 13.2±2.07, a-TNFR2 8.2±1.55; IFNγ and TNFα: control 22.46±2.96, a-TNFR2 9.5±2.04,(**j)** Branch length/cell index for TNFR2 neutralizations differentiation experiments: T3 only: control 1370.0±225.3, a-TNFR2 1344.0±51.36; IFNγ: control 737.8±124.9, a-TNFR2 758.2±37.1; TNFα: control 364.7±72.17, a-TNFR2 285.2±21.93; IFNγ and TNFα: control 1098.0±68.34, a-TNFR2 330.9±33.73,(**k)** Percentage of MHC-II+ cells out of NG2+ cells on day 3: IFNγ: control 33.71±1.12%, a-TNFR2 32.89±0.61%; IFNγ and TNFα: control 55.42±0.46%, a-TNFR2 54.51±1.2%; n=4 for each group,(**l)** Representative immunofluorescence analysis of MHC-II expression among OPCs upon TNFR2 neutralization and culture with either anti- or pro-inflammatory cytokines for 72h(scale bar = 40 μm),(**m)** CCL2 levels for TNFR2 neutralizations differentiation experiments: T3 only: control 290.1±44.59pg/mL, a-TNFR2 286.15±40.32pg/mL; IFNγ: control 835.24±33.9pg/mL, a-TNFR2 847.69±6.02pg/mL; TNFα: control 946.68±29.89pg/mL, a-TNFR2 999.7±82.29pg/mL; IFNγ and TNFα: control 1088.0±55.63pg/mL, a-TNFR2 1223.0±43.08pg/mL, n=4 for each group,(**n)** IL6 levels for TNFR2 neutralizations differentiation experiments: T3 only: control 10.69±2.67pg/mL, a-TNFR2 10.26±1.97pg/mL; IFNγ: control 20.67±0.66pg/mL, a-TNFR2 2053.0±0.95pg/mL; TNFα: control 12.82±0.66pg/mL, a-TNFR2 19.94±0.99pg/mL; IFNγ and TNFα: control 26.44±1.46pg/mL, a-TNFR2 32.08±2.22pg/mL, n=4 for each group,(**o)** IL10 levels for TNFR2 neutralizations differentiation experiments: T3 only: control 58.64±2.58pg/mL, a-TNFR2 58.04±3.91pg/mL; IFNγ: control 15.64±1.53pg/mL, a-TNFR2 16.52±1.66pg/mL; TNFα: control 24.23±0.95pg/mL, a-TNFR2 31.75±5.27pg/mL; IFNγ and TNFα: control 43.68±2.44pg/mL, a-TNFR2 50.6±2.97pg/mL, n=4 for each group. Significance was determined by two-way ANOVA analysis followed by Tukey’s multiple comparison analysis (p * ≤ 0.05, ** ≤ 0.01, *** ≤ 0.001, **** ≤ 0.0001, ns = not significant). Data are presented as means±SD. Each assay was repeated independently at least three times.

Upon stimulation with either TNFα or both TNFα and IFNγ simultaneously, anti-TNFR2 pre-treatment led to notably reduced morphological complexity, suggesting TNFR2’s role in modulating the immune activation of oligodendrocytes^11,27,37^. However, no such effect was observed in the T3 alone or IFNγ groups (Figure 7d-j).

These observations support that TNFR2 signaling plays a role both in OPC differentiation and immune activation.

#### 3.2. TNFR2 neutralization does not alter MHC-II expression in OPC

To elucidate TNFR2’s role in OPC immunomodulation, we assessed its effect on MHC-II expression in OPCs following simultaneous TNFα and IFNγ stimulation. OPCs were pre-treated with an anti-TNFR2 neutralizing antibody or PBS for 1h, followed by culture with MOG peptide, IFNγ, and TNFα.

Pre-treating OPCs with anti-TNFR2 resulted in a significant difference between exposure to IFNγ alone and combined exposure to TNFα and IFNγ (33.71±1.12% vs. 55.42±0.46%, p<0.0001, Figure 7k-l), mirroring the results in non-pre-treated OPCs (Figure 4b-c, Figure 7k-l). MHC-II expression was consistent between control and pre-treated OPCs across all conditions, and no significant difference was observed (Figure 4b-c, Figure 7k-l).

#### 3.3. TNFR2 neutralization increases the secretion of pro-inflammatory cytokines

Last, we aimed to investigate the role of TNFR2 in the reciprocal balance of immune function and the regenerative role of OPCs in the inflamed-CNS. Thus, we measured CCL2, IL6, and IL10 levels in the supernatants of OPCs’ differentiation cultures. OPCs were pre-treated with a neutralizing antibody directed to TNFR2 or PBS for 1h. Then, cultures were treated with T3^35^ and pro-inflammatory cytokines for seven days. Then, supernatants were collected and evaluated using ELISA.

In the TNFR2-neutralized group, OPCs simultaneously exposed to TNFα and IFNγ secreted higher levels of CCL2 compared to the control group(anti-TNFR2: 1223.0±43.08pg/mL; control: 1088.0±55.63pg/mL; p=0.0087). Elevated secretion levels were similarly observed for IL6 and IL10 in the anti-TNFR2 group (Figure 7m-o).

Together, these findings imply that TNFR2 plays a pivotal role in both the regenerative and immunomodulatory activities of OPCs within the inflamed-CNS.

## Discussion

In the current study, our objective was to delineate the influence of both pro- and anti-inflammatory environments on the immunological and differentiation functions of OPCs. We observed that exposure to an anti-inflammatory environment, represented by IL4 and IL10, resulted in decreased OPC differentiation. Interestingly, this effect was comparable to that induced by IFNγ, a predominant pro-inflammatory cytokine associated with MS. Moreover, exposure to IL4 and IL10 corresponded with reduced phagocytic activity, MHC-II expression, and cytokine secretion. Conversely, OPCs exposed to TNFα exhibited enhanced differentiation and robust immune functional capabilities. Those effects were attributed to signaling via TNFR2 and counteracted the detrimental effects of IFNγ on OPC differentiation. Our results support the hypothesis that the optimal functionality of OPCs requires a pro-regenerative inflammatory environment. Additionally, our results suggest the potential central role of TNFα and TNFR2 in orchestrating the equilibrium between the regenerative and immunological roles of OPCs within the inflamed-CNS.

For decades, the prevailing belief was that effective remyelination necessitates an anti-inflammatory environment. This belief has roots in some experimental findings. In an animal model examining stroke, IL4 administration was observed to enhance white matter integrity^21^. Additionally, oligodendrocyte differentiation was enhanced when exposed to conditioned media from microglia treated with anti-inflammatory cytokines(IL10,IL13)^22^. Nonetheless, it is essential to consider that this effect was compared with the effects of IFNγ and LPS, both detrimental to OPC differentiation. A critical observation is that these studies offered indirect insights and were conducted under varying conditions, potentially obscuring a direct understanding of the effects on OPCs.

Our data suggest that a conducive-permissive environment is essential for the effective functionality of the remyelinating cell. This may encompass pro-regenerative inflammatory elements, transitioning from a restrictive to a permissive environment. Inflammation’s beneficial role in promoting remyelination has been substantiated by extensive research. For instance, immune cells facilitate the removal of myelin debris, which contains proteins that inhibit OPC differentiation^41–43^. Moreover, remyelination occurs more frequently in active inflammatory MS lesions as compared to immunologically inactive plaques^44–47^. As confirmed by animal models of chronic demyelination, OPCs can only successfully remyelinate when acute inflammation is induced^48,49^. We demonstrated previously that exposure to CSF from rMS patients, which is characterized by heightened immune activity, led to enhanced OPCs’ differentiation, as compared to exposure to CSF from pMS patients, wherein the inflammatory milieu is relatively limited^11^. Additionally, pro-inflammatory milieu promotes OPC differentiation via interactions with activated microglia and lymphocyte cytokine secretion^50^. Also, experimental depletion of macrophages^51^, B or T cells^52,53^ leads to remyelination impairment. Together with our results, these suggest that remyelination depends on an appropriate immune response controlled in time, space, and intensity^54,55^.

There have been significant advances in elucidating OPCs’ immune-modulatory functions in various neurological disorders^15–17^. Following injury, OPCs are activated by inflammatory signals, leading them to secrete IL1β and CCL2. This facilitates their mobilization to the site of injury, subsequently augmenting the post-injury inflammatory environment^15,18,53,56,57^. The increased CCL2 levels upon TNFα exposure suggest a beneficial regenerative role for TNFα in the inflamed-CNS. Additionally, the elevated secreted IL6 levels in a pro-inflammatory context indicate its potential as a therapeutic target, where inhibiting IL6^58^ could support regeneration and neuroprotection. Furthermore, elevated IL10 levels following TNFα exposure highlight the necessary balance for CNS regeneration, suggesting a combined therapeutic approach with pro-inflammatory mediators to foster a conducive, regenerative inflammatory environment.

Immune-activated OPCs undergo morphological changes, including shortening and thickening processes, showing increased differentiation and myelination capability^11,27,37,38^. Consistently, OPCs exposed to TNFα exhibited higher differentiation and reduced morphological complexity. Further investigations have revealed that OPCs possess phagocytic abilities^17^ and have the capability to present antigens^11,17,19^. We demonstrated previously that OPC exposed to CSF from rMS patients exhibited augmented immune-modulatory functions(MHC-II expression, NFκB activation, cytokines secretion, T-cell activation and proliferation) compared to CSF from pMS patients where these functions appeared compromised^11^. This suggests that an increased inflammatory response of OPC is inversely associated with disease severity. Furthermore, merely pushing OPC to differentiate while suppressing their immune functions doesn’t adequately address the remyelination failure in chronic-MS^27^. These highlight the importance of the delicate balance between OPCs’ immune and pro-myelinating functions and underscore the need for a pro-inflammatory supporting environment for OPCs to execute their pro-myelinating and immune-modulatory roles.

In line with these concepts, our current findings spotlighted the potential protective and reparative effects of TNFα on OPC in the inflamed-CNS. TNFα exposure not only preserved the differentiation potential of OPCs but also sustained their immune-modulatory functions. Remarkably, TNFα appeared to counteract IFNγ’s detrimental effects on OPC differentiation, implying its potential beneficial role in remyelination in MS. These results align with recent insights regarding the role of TNFα and TNFR2 in OPC homeostasis. An EAE mice lacking oligodendroglial TNFR2 exhibited peripheral immune cell infiltration, increased demyelination, and impaired remyelination compared to wild-type^26^. Another research indicated that TNFR2 modulates OPCs’ pro-inflammatory phenotype^25^. The absence of TNFR2 exacerbates the immune-modulatory and inflammatory response in OPCs post-inflammatory induction, consequently reducing their proliferation and differentiation potential^25^. An EAE model with dual modulation of TNFR1 and TNFR2 yielded promising results, effectively mitigating symptoms while decreasing demyelination and axonal degeneration^59^. The pivotal function of TNFα in maintaining a CNS regenerative milieu is further underscored by instances where patients treated with anti-TNF agents developed demyelinating syndromes^60^. These findings highlight the significance of immune signaling in determining OPC functions and indicate the promising therapeutic potential of targeting TNFα and TNFR2 pathways in MS.

In conclusion, our study sheds lights on the nuanced interplay between pro- and anti-inflammatory environments and the subsequent impact on OPCs’ immunological and differentiation functionalities. Our findings point towards the essentiality of a pro-regenerative inflammatory environment, emphasizing the pivotal role of TNFα and TNFR2 in harmonizing the balance between the regenerative and immunological tasks of OPCs in the inflamed-CNS. These findings may pave the way for innovative therapies that encapsulate the full spectrum of OPC functions in MS.

